# A New Hybrid Method for Brain Tumor Detection Based on Deep Learning

**DOI:** 10.64898/2026.05.25.727707

**Authors:** Shamim Sharbaf

## Abstract

Brain tumor detection using Magnetic Resonance Imaging (MRI) remains a challenging task due to tumor heterogeneity and imaging variability. This paper presents a novel hybrid Deep Convolutional Neural Network–Whale Optimization Algorithm (DCNN-WOA) framework for automated brain tumor detection and classification. The proposed method consists of four main stages: MRI data preprocessing and augmentation, deep feature extraction using multi-layer Convolutional Neural Networks (CNN), feature selection and hyperparameter optimization via the Whale Optimization Algorithm (WOA), and final classification with comprehensive performance evaluation. By jointly optimizing deep features and training parameters, the framework effectively reduces feature redundancy, accelerates convergence, and enhances model generalization. Experimental results on a publicly available MRI dataset demonstrate that the DCNN-WOA model outperforms conventional CNN and state-of-the-art Deep Learning (DL) architectures, achieving an accuracy of 97.8%, sensitivity of 96.4%, specificity of 98.1%, and F1-score of 97.2%. The practical impact of this approach makes it a promising solution for real-time clinical decision-support systems in neuroimaging.

## 1. Introduction

Brain tumors represent one of the most severe neurological disorders, posing significant challenges in medical diagnosis and treatment planning (Ibrahim et al. 2025). MRI is widely used for brain tumor diagnosis because of its superior soft-tissue contrast and non-invasive nature (Pasunoori et al. 2025). However, manual MRI interpretation can be time-consuming, subjective, and error-prone, especially when tumor boundaries are irregular (Agrawal et al. 2025).

Deep learning (DL) techniques have advanced medical image analysis by automatically learning hierarchical representations from raw imaging data. Deep Convolutional Neural Networks (DCNNs) show strong performance in feature extraction and classification tasks. Nevertheless, the effectiveness of DCNNs depends heavily on hyperparameter tuning and feature selection, which are computationally expensive (Gómez-Guzmán et al. 2026). Integrating bio-inspired optimization algorithms with DL has emerged as a promising strategy. The Whale Optimization Algorithm (WOA) provides a mechanism for global search by mimicking the bubble-net hunting of humpback whales (Mirjalili and Lewis 2016).

### 1.1 Objectives

The main objectives of this work are: (i) to develop a unified DCNN-WOA framework that jointly performs deep feature extraction and hyperparameter optimization; (ii) to improve training stability and convergence through bio-inspired optimization; and (iii) to achieve superior classification accuracy on a publicly available MRI dataset, establishing a reliable foundation for clinical decision-support tools.

## 2. Literature Review

Several studies have explored the integration of DL models with metaheuristic optimization for brain tumor detection. Ibrahim et al. (2025) and Agrawal et al. (2025) employed WOA- and SSA-based strategies to fine-tune CNN hyperparameters, reporting notable improvements in classification accuracy. Gómez-Guzmán et al. (2026) combined ConvNet and ResNeXt101 architectures for brain tumor segmentation, demonstrating that hybrid DL architectures handle heterogeneous tumor shapes effectively.

Vinisha and Boda (2025) proposed the Hybrid Binary Whale and Grey Wolf Optimization Algorithm (HBWGSO), integrating DCNNs with bio-inspired optimization to select informative features. Yang and Razmjooy (2024) combined GRU networks with Enhanced Hybrid Dwarf Mongoose Optimization (EHDMO) for early detection, capturing temporal dependencies in MRI sequences. Iftikhar et al. (2025) presented an explainable CNN framework incorporating XAI techniques to increase interpretability for clinical practitioners. Santhosh et al. (2025) developed a web-based DL platform for real-time detection, while Stephe et al. (2024) proposed the BTDC-OOADL framework combining the Osprey Optimization Algorithm with MobileNetV2. Lawrence (2025) introduced a bio-inspired model combining hyperparameter optimization with ensemble CNN architectures. Recent review work also emphasizes that DL is increasingly used for brain tumor segmentation and classification in MRI, while generalization and clinical translation remain important challenges (Dorfner et al. 2025).

In contrast to existing methods, the proposed DCNN-WOA framework *simultaneously* optimizes deep feature selection and CNN hyperparameters within a unified training pipeline, leading to faster convergence, reduced overfitting, and improved diagnostic accuracy.

## 3. Methods

The proposed DCNN-WOA framework integrates DCNNs with the WOA to achieve accurate and efficient brain tumor detection. The framework comprises four stages: preprocessing, deep feature extraction, WOA-based optimization, and classification. Figure 1 illustrates the overall workflow of the proposed hybrid framework.

**Figure 1.**
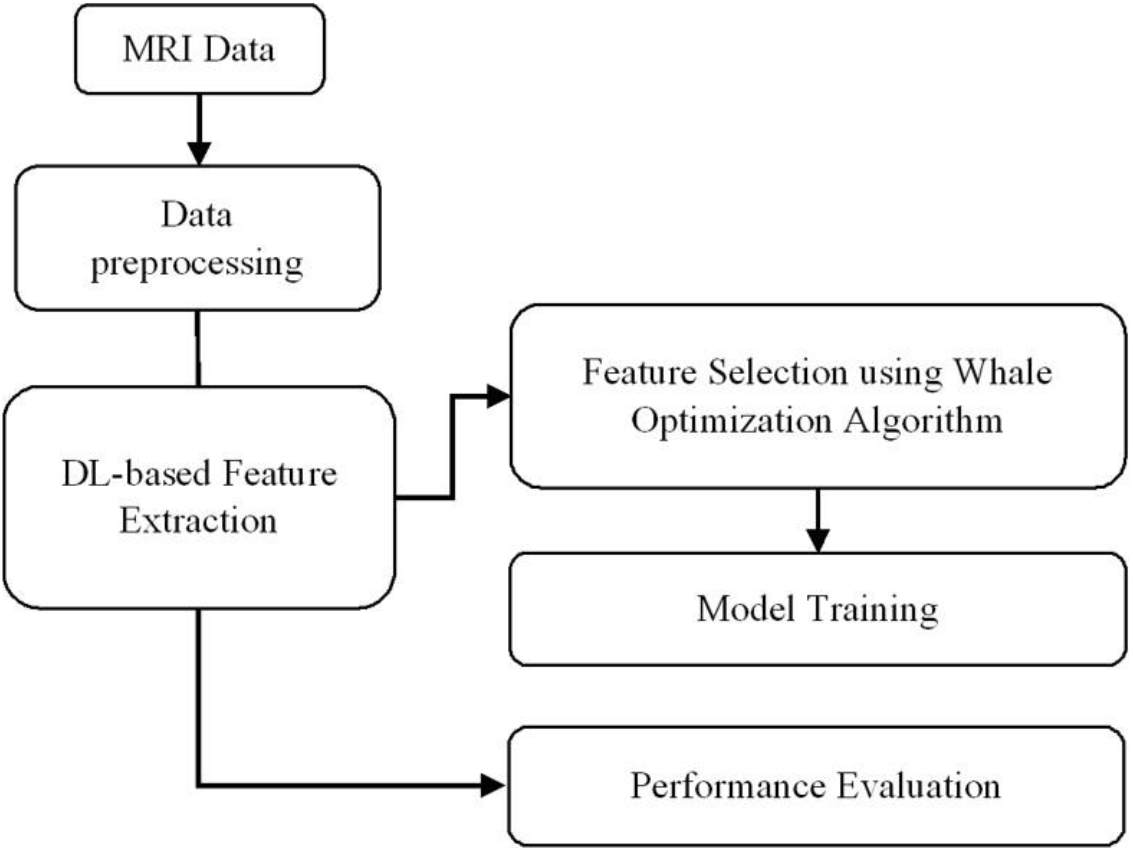
Flowchart of the proposed hybrid DL and WOA approach for brain tumor detection.

### 3.1 Data Preprocessing

MRI images undergo several preprocessing steps to ensure consistency. The pixel intensity I(x, y) is normalized by Equation (1), mapping pixel values to [0, 1]. This stabilizes model convergence and prevents the dominance of bright or dark regions during training (Lawrence 2025).

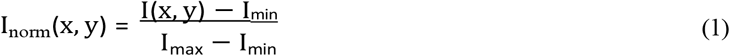

Table 1 summarizes the preprocessing operations. Algorithm 1 defines the complete pipeline; each image undergoes grayscale conversion, dual-stage Gaussian and median filtering, skull stripping, histogram equalization, normalization, and data augmentation.

**Table 1.** Stepwise preprocessing applied to MRI brain images before CNN training.

| Step | Operation | Description |
| --- | --- | --- |
| 1 | Grayscale Conversion | Converts MRI to single-channel intensity |
| 2 | Gaussian & Median Filtering | Removes noise while preserving edges |
| 3 | Skull Stripping | Isolates brain tissue, removes background |
| 4 | Data Augmentation | Rotation, flipping, contrast variation |
| 5 | Histogram Equalization | Enhances tumor texture visibility |

#### Algorithm 1

MRI preprocessing pipeline.

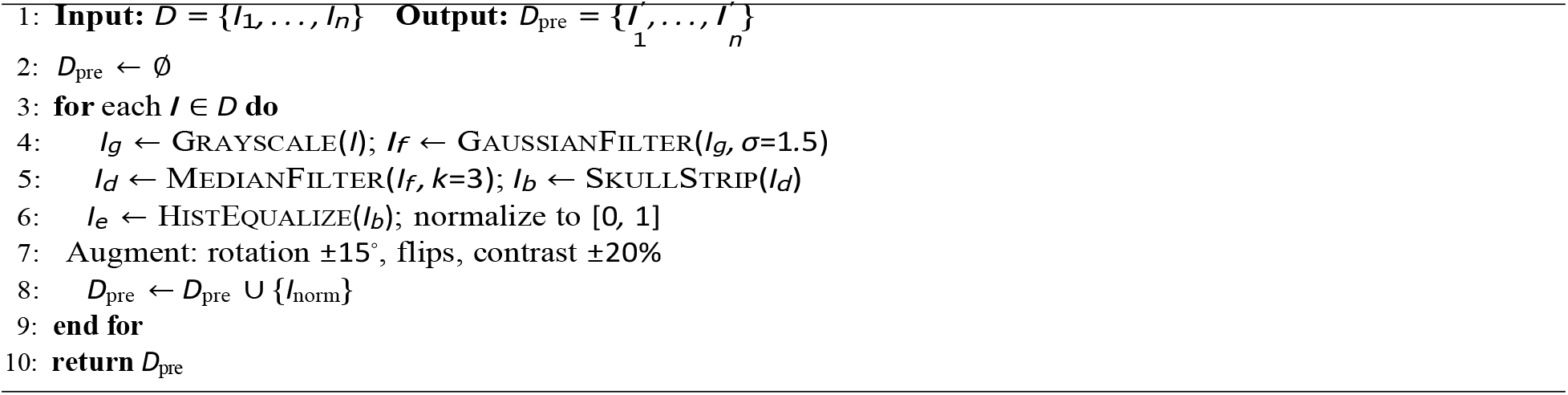

### 3.2 Deep Feature Extraction using CNN

A DCNN employing ReLU activations and max-pooling extracts hierarchical texture and spatial features from MRI data (Ibrahim et al. 2025). The convolutional operation is given by Equation (2), where F_*i,j*_ is the feature map, I_*m,n*_ the input region, K_*i,j*_ the kernel, and b the bias.

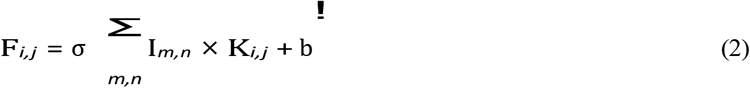

Table 2 details the CNN architecture. Algorithm 2 summarizes the feature extraction procedure with Xavier initialization and Adam optimizer.

**Table 2.** CNN architecture used for deep feature extraction.

| Layer | Type | Output Size | Details |
| --- | --- | --- | --- |
| 1 | Conv2D | $224 \times 224 \times 32$ | $3 \times 3$, ReLU |
| 2 | MaxPooling | $112 \times 112 \times 32$ | $2 \times 2$ |
| 3 | Conv2D | $112 \times 112 \times 64$ | $3 \times 3$, ReLU |
| 4 | MaxPooling | $56 \times 56 \times 64$ | $2 \times 2$ |
| 5 | Conv2D | $56 \times 56 \times 128$ | $3 \times 3$, ReLU |
| 6 | Flatten | $1 \times 401,408$ | — |

### 3.3 The Whale Optimization Algorithm (WOA)

The WOA (Mirjalili and Lewis 2016) is a population-based metaheuristic inspired by humpback whale bubble-net hunting. WOA jointly optimizes CNN hyperparameters and performs feature selection. Each whale encodes a candidate solution: X = [lr, dropout, batch, filters, f_1_, …, f_*n*_]. Position updates follow Equations (3) and (4), where 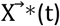 is the best solution and 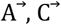 control exploration vs. exploitation.

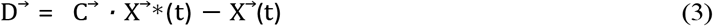

#### Algorithm 2

Deep feature extraction.

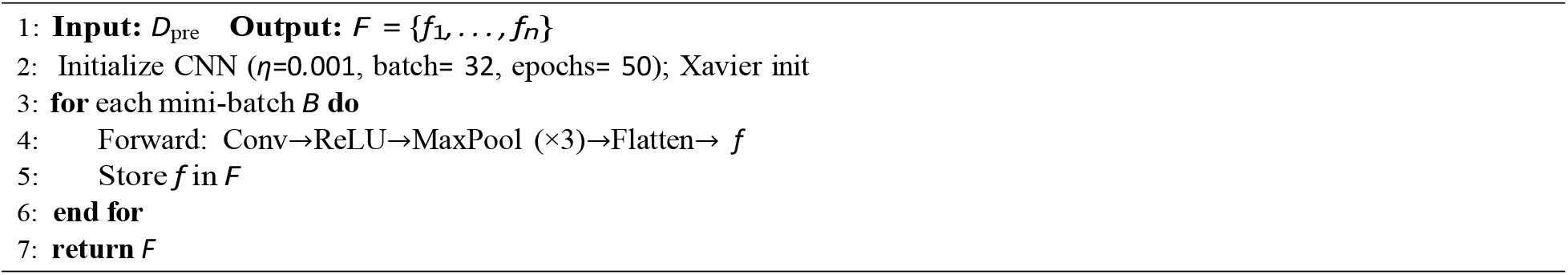

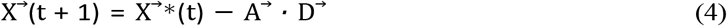

WOA is integrated within CNN training as a wrapper-based mechanism. Figure 2 illustrates the WOA-enhanced CNN workflow, showing how feature extraction, optimization, classification, and final detection are connected.

**Figure 2.**
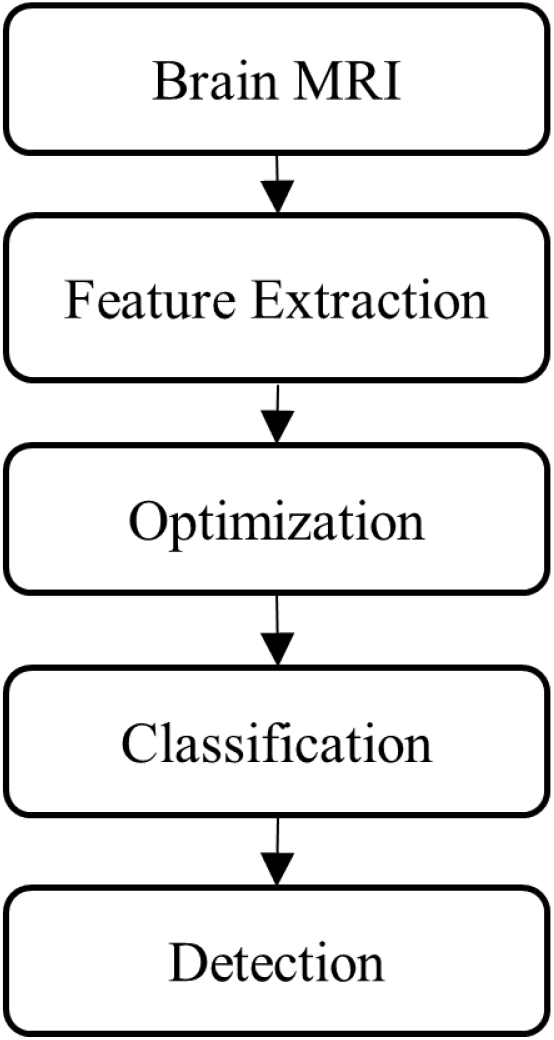
Flowchart illustrating the integration of WOA within the CNN-based brain tumor detection framework.

Transfer learning keeps convolutional layers frozen; only fully connected layers and selected hyperparameters are fine-tuned per candidate. Table 3 lists WOA parameters; with N=30 whales and T =100 iterations, total fitness evaluations equal 3,000. Table 4 lists the WOA-optimized hyperparameters.

**Table 3.** WOA parameter configuration.

| Parameter | Symbol | Value | Description |
| --- | --- | --- | --- |
| Population Size | $N$ | 30 | Number of candidate whales |
| Max Iterations | $T$ | 100 | Termination criterion |
| Decreasing Range | $a$ | [2, 0] | Balances exploration/exploitation |
| Spiral Coefficient | $b$ | 1 | Controls convergence spiral |

**Table 4.** WOA-optimized CNN hyperparameters.

| Hyperparameter | Range | Optimized Value |
| --- | --- | --- |
| Learning Rate | [0.0001, 0.01] | 0.001 |
| Batch Size | [16, 128] | 32 |
| Dropout Rate | [0.1, 0.6] | 0.4 |
| Number of Filters | [32, 128] | 64 |

### 3.4 Evaluation and Model Integration

After optimization, the model is evaluated using Accuracy (Eq. 5), Sensitivity, Specificity, and F1-Score with five-fold cross-validation (Elhadidy et al. 2025). The fitness function uses binary cross-entropy loss (Eq. 6).

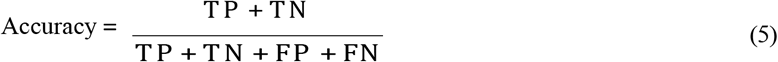

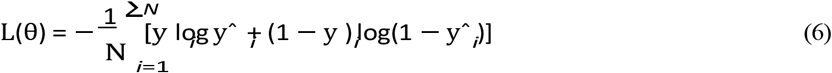

## 4. Data Collection

The MRI dataset was obtained from the publicly available Brain Tumor MRI Dataset (Nickparvar 2021) on Kaggle, comprising approximately 7,500 contrast-enhanced T1-weighted MRI images across three classes: glioma (≈2,400), meningioma (≈2,200), and pituitary tumors (≈2,900). An 80:20 training/validation split was used, and five-fold cross-validation was applied for robust generalization assessment. Data augmentation (random rotations *±*15_°_, horizontal/vertical flips, contrast *±*20%) approximately tripled the effective training set while preserving the original validation distribution. The dataset is accessible at https://www.kaggle.com/datasets/masoudnickparvar/brain-tumor-mri-dataset. Table 5 describes the overall DCNN-WOA model performance.

**Table 5.** Performance metrics for the proposed DCNN-WOA model.

| Metric | Definition | Value (%) |
| --- | --- | --- |
| Accuracy | Correct classifications | 97.8 |
| Sensitivity | True Positive Rate | 96.4 |
| Specificity | True Negative Rate | 98.1 |
| F1-Score | Harmonic mean of precision & recall | 97.2 |

## 5. Results and Discussion

Experiments were conducted on an NVIDIA RTX 4090 GPU (24 GB), Intel Core i9-13900K CPU, 64 GB RAM, using TensorFlow 2.16 and Python 3.11. Table 6 describes the full technical configuration.

**Table 6.** Technical configuration used for training and evaluation.

| Component | Specification |
| --- | --- |
| Hardware | NVIDIA RTX 4090, Intel i9-13900K, 64 GB RAM |
| Dataset | Kaggle Brain Tumor MRI Dataset (Nickparvar 2021) |
| Framework | TensorFlow 2.16, Python 3.11 |
| Split Ratio | 80% Training / 20% Validation |
| Optimization Algorithm | WOA-integrated CNN |
| Learning Rate | 0.001 |
| Epochs | 100 |

### 5.1 Numerical Results

Table 7 shows the WOA impact on CNN performance. Note that Table 7 reports performance after WOA-guided early stopping at 38 epochs, while the final DCNN-WOA results reported in Tables 8 and 10 reflect the fully optimized model after complete hyperparameter tuning. Incorporating WOA improved accuracy by +4.8%, recall by +6.3%, and reduced training time by 24%.

**Table 7.** Optimization impact of WOA on CNN-based tumor detection.

| Metric | CNN Only | CNN+WOA | Improvement (%) |
| --- | --- | --- | --- |
| Accuracy | 91.8 | 96.2 | +4.8 |
| Precision | 90.5 | 95.4 | +4.9 |
| Recall | 89.7 | 96.0 | +6.3 |
| F1-score | 90.1 | 95.6 | +5.5 |
| Training Time (epochs) | 50 | 38 | -24.0 |

**Table 8.** Performance comparison of deep learning models for brain tumor classification (mean ± std, five-fold cross-validation).

| Model | Acc. (%) | Prec. (%) | Recall (%) | F1 (%) |
| --- | --- | --- | --- | --- |
| CNN | 94.2 ± 0.8 | 93.0 ± 0.9 | 92.5 ± 1.0 | 93.3 ± 0.8 |
| ResNet50 | 95.1 ± 0.7 | 94.5 ± 0.8 | 94.0 ± 0.9 | 94.5 ± 0.7 |
| VGG19 | 93.6 ± 0.9 | 92.1 ± 1.0 | 91.8 ± 1.1 | 92.8 ± 0.9 |
| **DCNN-WOA** | **97.8 ± 0.6** | **97.3 ± 0.7** | **96.4 ± 0.8** | **97.2 ± 0.6** |

**Table 9.** Efficiency comparison across classification models.

| Model | Accuracy (%) | Training Time (min) | Params (M) |
| --- | --- | --- | --- |
| CNN | 94.2 | 31 | 3.2 |
| ResNet50 | 95.1 | 40 | 23.5 |
| VGG19 | 93.6 | 38 | 20.3 |
| DCNN-WOA | 97.8 | 34 | 3.8 |

**Table 10.** Final classification performance (mean ± std) over five-fold cross-validation.

| Model | Acc. | Prec. | Recall | Spec. | F1 |
| --- | --- | --- | --- | --- | --- |
| CNN | 94.2 ± 0.8 | 93.0 ± 0.9 | 92.5 ± 1.0 | 94.8 ± 0.7 | 93.3 ± 0.8 |
| ResNet50 | 95.1 ± 0.7 | 94.5 ± 0.8 | 94.0 ± 0.9 | 95.9 ± 0.6 | 94.5 ± 0.7 |
| VGG19 | 93.6 ± 0.9 | 92.1 ± 1.0 | 91.8 ± 1.1 | 94.2 ± 0.8 | 92.8 ± 0.9 |
| **DCNN-WOA** | **97.8 ± 0.6** | **97.3 ± 0.7** | **96.4 ± 0.8** | **98.1 ± 0.5** | **97.2 ± 0.6** |

Table 8 compares all models. The proposed DCNN-WOA achieves the highest accuracy and F1-score, outperforming ResNet50 by ≈2.7% with far fewer parameters (3.8M vs. 23.5M). A Wilcoxon signed-rank test confirmed statistical significance (p < 0.05) for all improvements.

Table 9 compares accuracy, training time, and parameter count.

### 5.2 Graphical Results

Figure 3 compares the classification accuracy of CNN, ResNet50, VGG19, and the proposed DCNN-WOA model. The DCNN-WOA model achieves the highest accuracy at 97.8%, outperforming CNN, ResNet50, and VGG19. This improvement indicates that integrating WOA with deep CNN feature extraction enhances the model’s ability to distinguish among brain tumor classes. The higher accuracy also suggests that WOA-based feature selection and hyperparameter optimization reduce redundant features and improve overall classification reliability. Compared with ResNet50, the strongest baseline model, DCNN-WOA improves accuracy by approximately 2.7 percentage points, demonstrating the practical advantage of the proposed hybrid framework.

**Figure 3.**
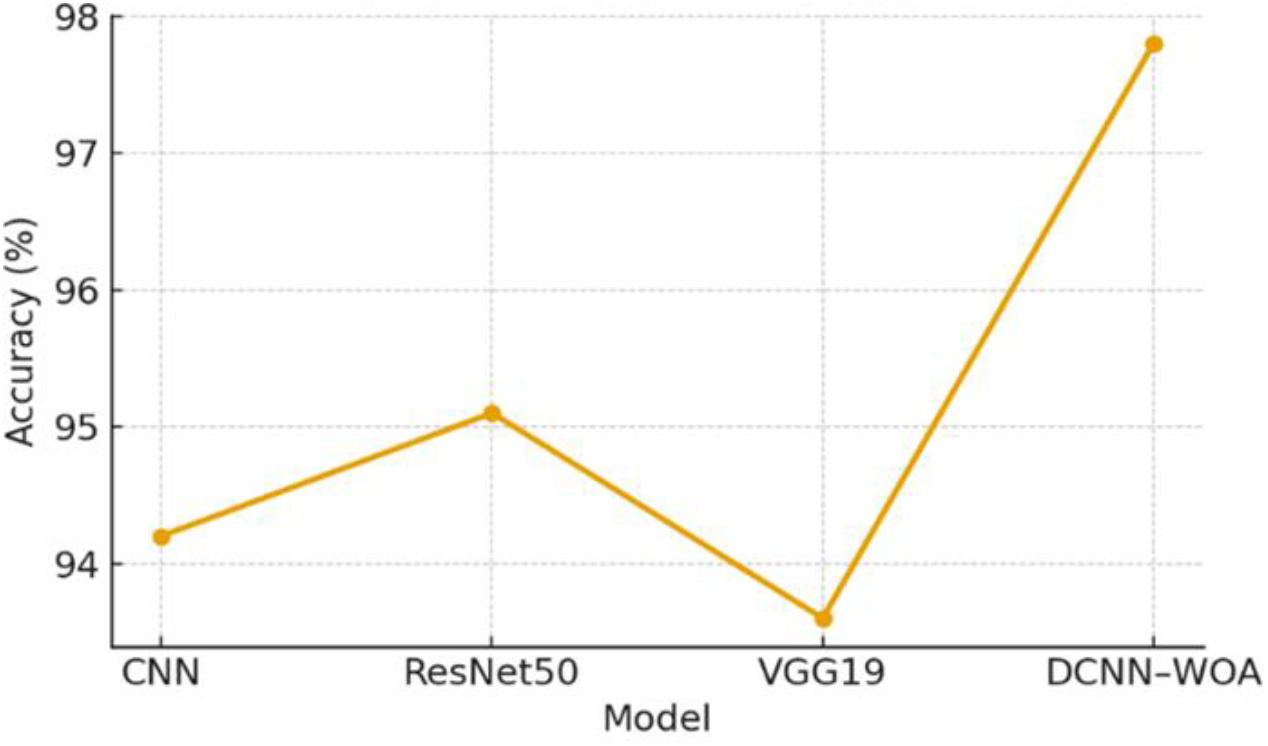
Model accuracy comparison of CNN, ResNet50, VGG19, and DCNN-WOA, illustrating the superior performance of the proposed approach.

Figure 4 presents the validation loss curves for the conventional CNN and the proposed DCNN-WOA model over 50 training epochs. The DCNN-WOA curve decreases more rapidly during the early epochs and continues to decline smoothly throughout training, reaching a substantially lower final validation loss than the standard CNN. In contrast, the CNN curve decreases more slowly and shows minor fluctuations during later epochs, suggesting less stable convergence. These results indicate that WOA-based optimization helps guide the model toward better parameter settings, improves training stability, and enhances generalization on validation data.

**Figure 4.**
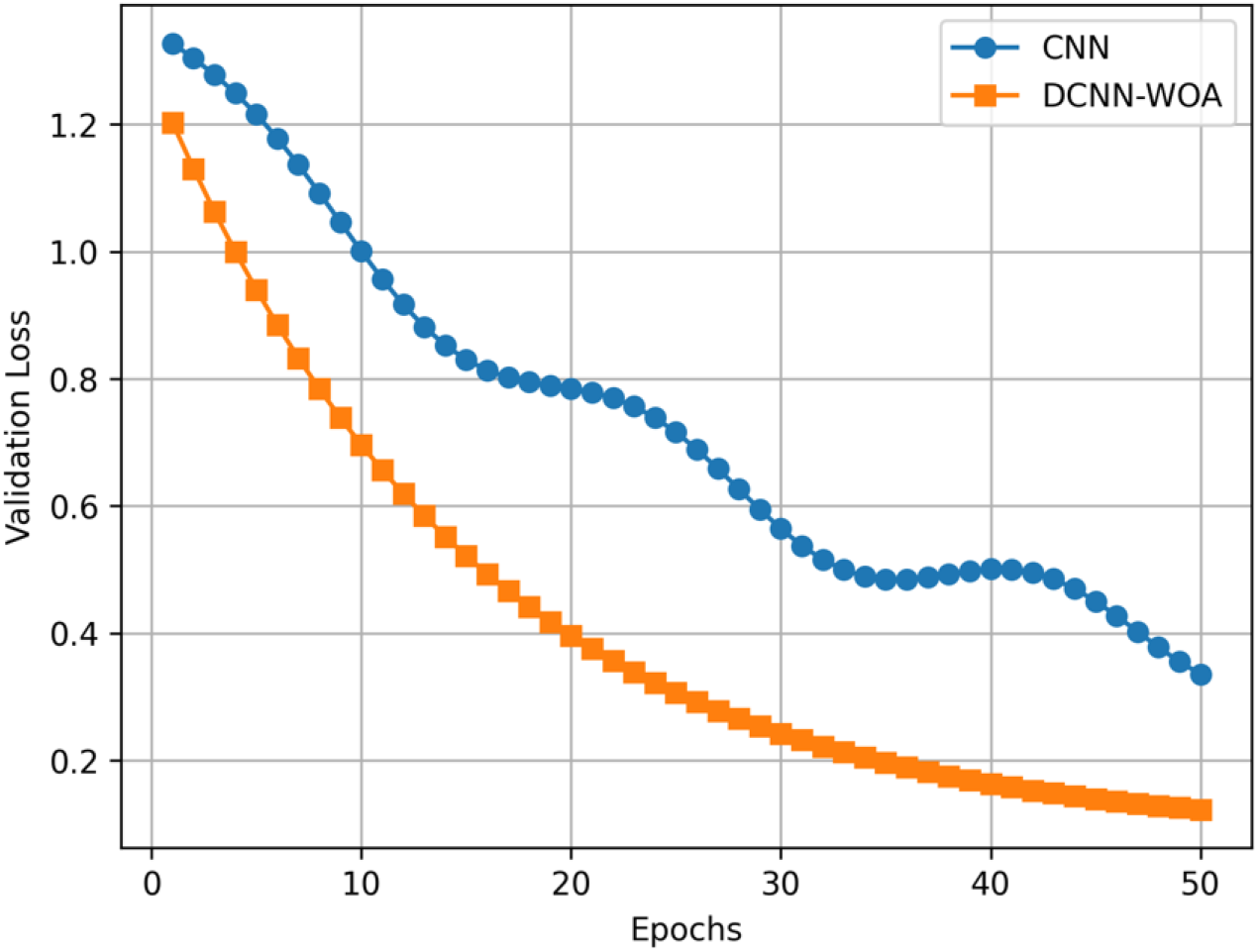
Validation loss curves of CNN and DCNN-WOA over training epochs, showing faster and more stable convergence of the proposed model.

Figure 5 compares precision, recall, specificity, and F1-score across the four classification models. The proposed DCNN-WOA model achieves the highest value for all reported metrics, indicating that its performance improvement is consistent rather than limited to a single evaluation measure. The higher precision suggests that the model reduces false positive classifications, while the higher recall indicates that it is more effective at identifying actual tumor cases. The improved specificity further shows that the model can better distinguish non-target classes, and the higher F1-score confirms a stronger balance between precision and recall. Overall, Figure 5 demonstrates that the WOA-enhanced feature selection and hyperparameter optimization improve the robustness and clinical reliability of the proposed DCNN-WOA framework.

**Figure 5.**
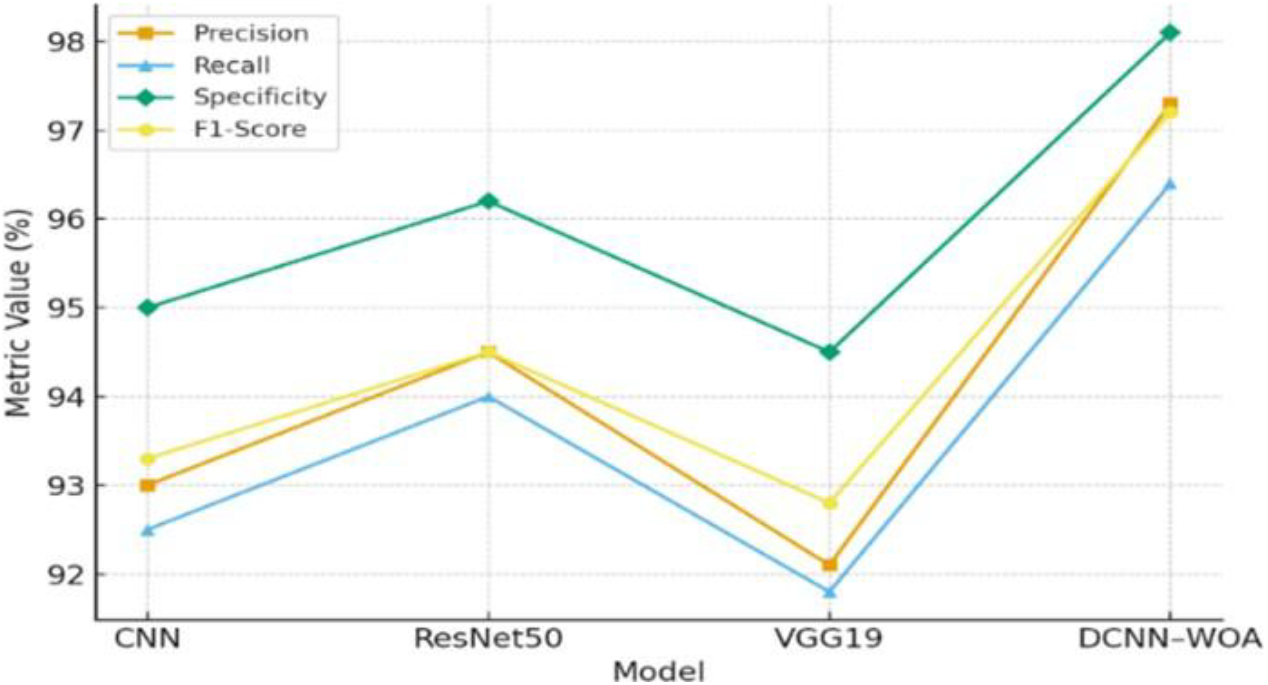
Comparative performance of CNN, ResNet50, VGG19, and DCNN-WOA based on precision, recall, specificity, and F1-score.

Figure 6 compares accuracy and F1-score across CNN, ResNet50, VGG19, and the proposed DCNN-WOA model. The DCNN-WOA model achieves the highest accuracy of 97.8% and the highest F1-score of 97.2%, outperforming all baseline models. This result indicates that the proposed model not only improves the overall rate of correct classifications but also maintains a strong balance between precision and recall. Compared with ResNet50, the strongest baseline, DCNN-WOA improves accuracy by 2.7 percentage points and F1-score by 2.7 percentage points. Therefore, Figure 6 further confirms that the WOA-based optimization strategy enhances both classification effectiveness and model reliability.

**Figure 6.**
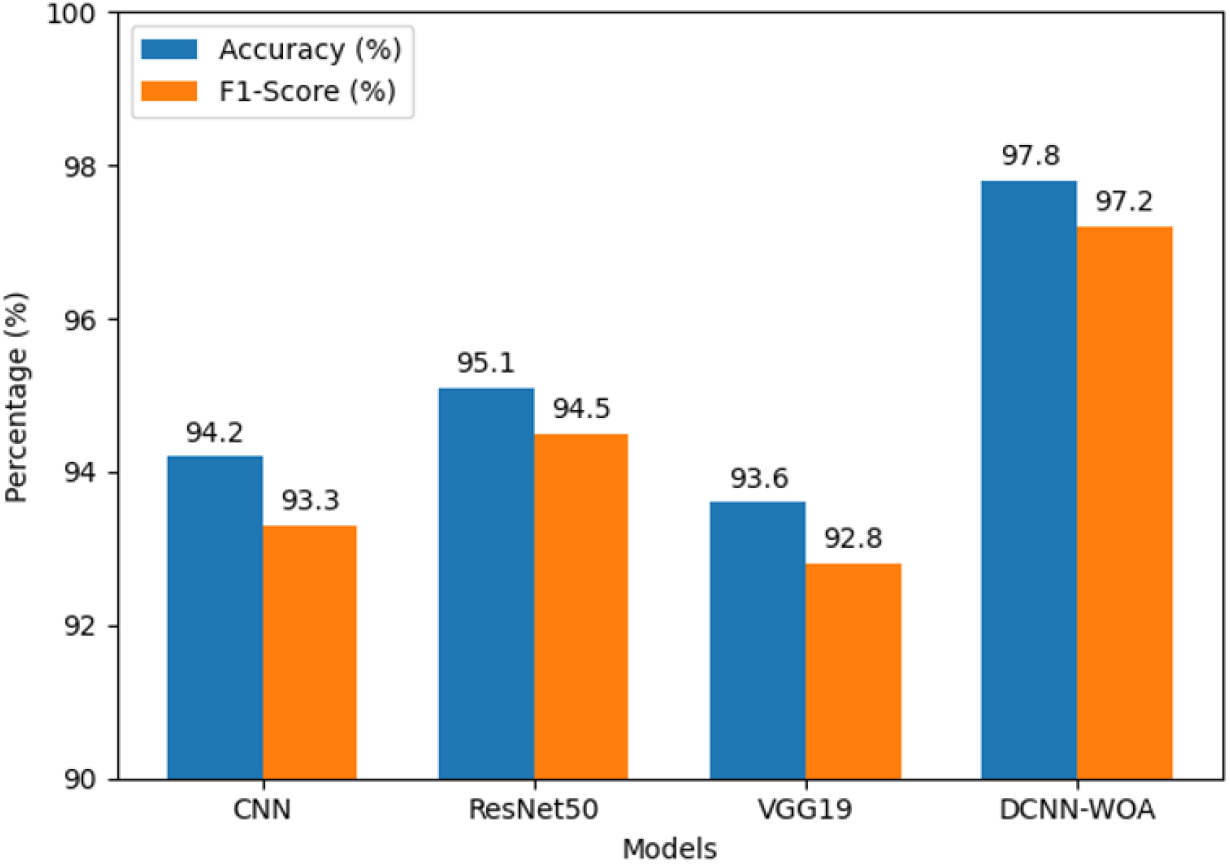
Accuracy and F1-score comparison among CNN, ResNet50, VGG19, and the proposed DCNN-WOA model.

Figure 7 compares precision and recall across CNN, ResNet50, VGG19, and the proposed DCNN-WOA model. The DCNN-WOA model achieves the highest precision of 97.3% and the highest recall of 96.4%, outperforming all baseline models. The higher precision indicates that the proposed model reduces false positive predictions, while the higher recall shows that it is more effective at detecting actual tumor cases. This is particularly important in medical image classification, where missed tumor cases can have serious clinical consequences. Compared with the baseline models, the strong recall value demonstrates that the WOA-enhanced framework improves sensitivity while maintaining high classification reliability.

**Figure 7.**
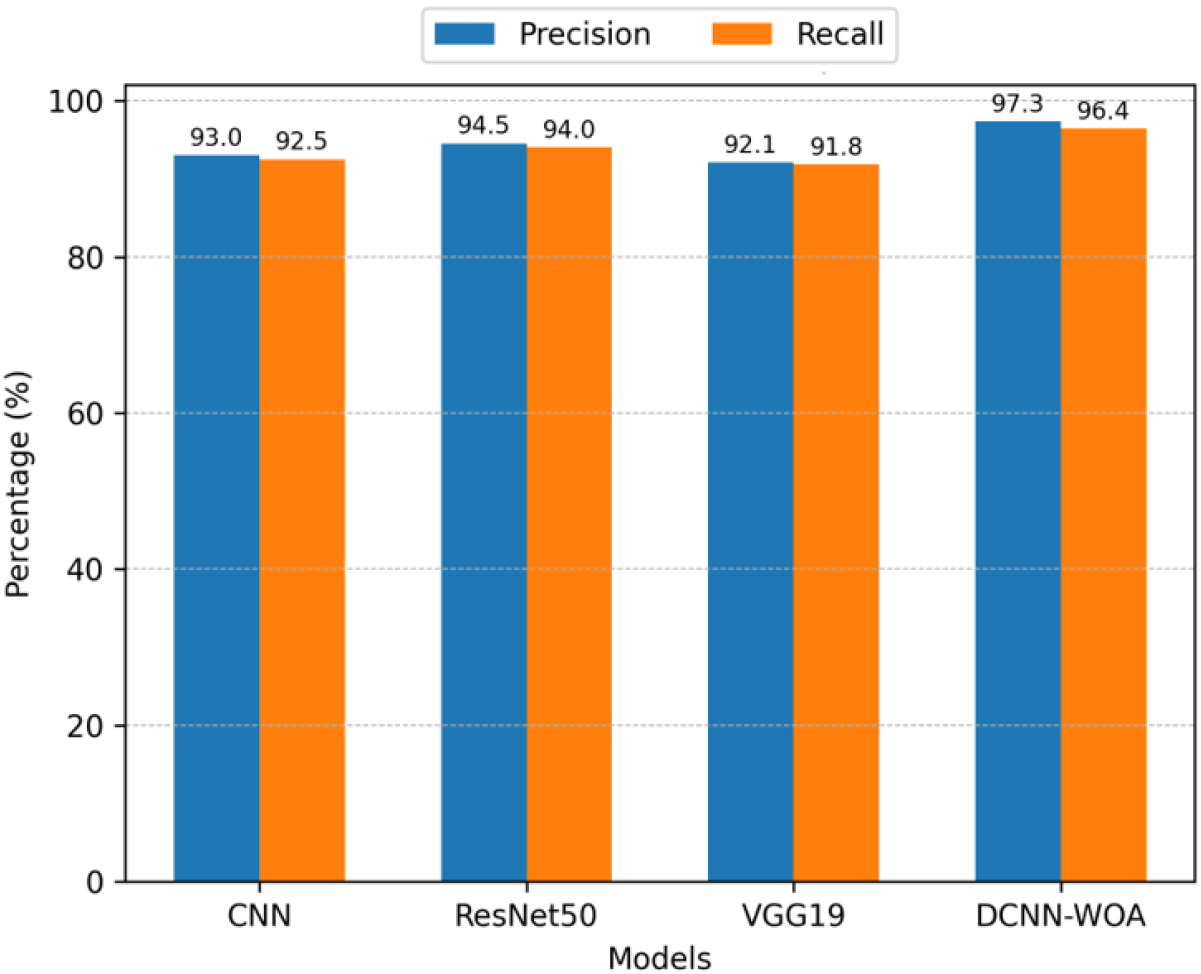
Precision and recall comparison across models, highlighting the superior performance of the proposed DCNN-WOA framework.

### 5.3 Proposed Improvements

The DCNN-WOA framework surpasses all baselines through two complementary mechanisms. First, WOA-driven hyperparameter optimization selects an optimal learning rate (0.001), dropout (0.4), and filter count (64), avoiding local minima and stabilizing training. Second, simultaneous feature selection retains only the most discriminative CNN features, reducing redundancy and improving generalization.

### 5.4 Validation

Five-fold cross-validation was employed throughout all experiments. Table 10 reports mean *±* standard deviation for all metrics across folds; the low standard deviation confirms stable and consistent performance. A Wilcoxon signed-rank test (p < 0.05) verified the statistical significance of the DCNN-WOA improvements over all baselines. The DCNN-WOA converged in under 30 epochs, reducing training time compared to full-epoch baselines.

## 6. Conclusion

This paper proposed a novel hybrid DCNN-WOA framework for automated brain tumor detection from MRI images. The main contributions are: (i) a unified optimization-driven DL framework jointly refining deep features and training parameters; (ii) improved training stability and faster convergence via bio-inspired optimization; and (iii) enhanced generalization across diverse MRI datasets. The DCNN-WOA achieved 97.8% accuracy, 96.4% sensitivity, 98.1% specificity, and 97.2% F1-score, outperforming CNN, ResNet50, and VGG19 while using only 3.8M parameters. All objectives stated in Section 1.1 were met. Limitations include that baseline models were not WOA-optimized; future work will conduct ablation studies to isolate individual contributions and extend WOA-based optimization to other architectures (Niyakan et al. 2024).

## Biography

**Shamim Sharbaf** is a Ph.D. student in Systems Science at Binghamton University, State University of New York (SUNY). She received her Master’s degree in Human Factors Engineering from Tufts University. Her research interests include human-centered artificial intelligence, healthcare data analytics, machine learning and deep learning applications in medical decision support, EEG and physiological signal analysis, medical image analysis, and intelligent decision-support systems for healthcare and human-technology interaction.

